# Similar network compositions, but distinct neural dynamics underlying belief updating in environments with and without explicit outcomes

**DOI:** 10.1101/794669

**Authors:** Vincenzo G. Fiore, Xiaosi Gu

## Abstract

Classic decision theories, such as reinforcement learning, typically require the presence of explicit outcomes for learning and belief updating. However, ecological environments are often opaque and explicit feedback, such as those based on values, might not be immediately accessible. It remains unclear whether the neural dynamics underlying belief updating in absence of outcomes differ from those responsible for decision-making based on accessible outcomes. Here, we investigated this question in healthy humans (n=28) using Bayesian modeling and two multi-option fMRI tasks, one with and one without immediate outcome. Model-based fMRI analysis revealed two opposing networks encoding belief updating regardless of the presence of immediate outcomes. A “confidence-building” network including the hippocampus, amygdala, and medial prefrontal cortex (mPFC) became more active as beliefs about action-outcome probabilities were confirmed by newly acquired information. Meanwhile, an “uncertainty-building” network including the anterior insular (AIC), dorsal anterior cingulate (dACC), and dorsolateral prefrontal (dlPFC) cortices became more active as new evidence conflicted with action-outcome estimates. Interestingly, dynamic causal modeling revealed that the confidence network was driven either by the hippocampus when outcomes were not available, or by the mPFC and amygdala when value-based outcomes were immediately accessible. Convsersely, the AIC always drove the activities of dACC and dlPFC, under the modulation of increasing uncertainty, independent of outcome availability. These findings reveal similar network compositions but distinct neural dynamics underlying belief updating in changing environments with and without explicit outcomes, highlighting an asymmetric relationship between decision confidence and uncertainty computation across levels of analysis.

**Highlights:**

- We investigated belief updating in two tasks, with and without explicit feedback.
- Model-based fMRI analysis revealed similar neural responses across tasks.
- The anterior insula drove an uncertainty-encoding network, across tasks.
- The anterior hippocampus drove a confidence-encoding network, w/o feedbacks.
- The medial PFC and amygdala drove a confidence-encoding network, with feedbacks.

## Introduction

In classic decision-making theories, the presence of explicit outcomes (e.g. based on values) after a choice selection is considered a crucial source of information for belief updating and behavioral adaptability, e.g. by triggering prediction error signals (Glimcher, 2011; Rangel, Camerer, & Montague, 2008). This line of research has yielded fruitful results, most prominently in the identification of the neurocomputational mechanisms underlying reinforcement learning (Dabney, et al., 2020; Schultz, Dayan, & Montague, 1997) and belief updates in changing environments (Behrens, Woolrich, Walton, & Rushworth, 2007; McGuire, Nassar, Gold, & Kable, 2014; Soltani & Izquierdo, 2019). In real life, however, decisions are often made in opaque environments where outcomes can be sporadic or temporarily inaccessible. Despite this opacity, we can still form and update beliefs about how likely our chosen actions are to deliver what we need or want, based on other sources of information (Ma & Jazayeri, 2014; Pouget, Beck, Ma, & Latham, 2013). For instance, previous studies have reported that, in probabilistic environments without explicit outcomes, the anterior hippocampus monitors the entropy in sequences of visual or auditory events in order to generate expectations regarding future stimuli and guide behavior accordingly (Harrison, Duggins, & Friston, 2006; Krug, et al., 2014; Strange, Duggins, Penny, Dolan, & Friston, 2005; Tobia, Iacovella, & Hasson, 2012). However, it is yet to be determined whether action-outcome belief updating under different conditions of explicit feedback access relies on different neural patterns and network dynamics.

In Bayesian terms, beliefs represent – as probability distributions – the estimated likelihood that one’s action will produce one or more known outcomes (Fleming & Daw, 2017; Kording & Wolpert, 2006; Payzan-LeNestour & Bossaerts, 2011). These probabilistic distributions are taken into account in decision-making (Orban & Wolpert, 2011; Sanders, Hangya, & Kepecs, 2016) and are encoded by the activity of neuronal populations (Ma, Beck, Latham, & Pouget, 2006; Pouget, et al., 2013; Rich, Cazettes, Wang, Pena, & Fischer, 2015). Under conditions in which sensory stimuli are unambiguous and reliable (Ma & Jazayeri, 2014), confirming accumulating evidence leads to narrow distributions and precise action-outcome beliefs, whereas conflicting information leads to wide distributions and imprecision (Meyniel, Schlunegger, & Dehaene, 2015; Meyniel, Sigman, & Mainen, 2015; Payzan-LeNestour, Dunne, Bossaerts, & O’Doherty, 2013; Pouget, Drugowitsch, & Kepecs, 2016). Thus, it has been proposed that “decision confidence” and “decision uncertainty” can be operationally considered as complementary probabilities (Adler & Ma, 2018; Atiya, et al., 2020; Meyniel, Schlunegger, et al., 2015; Meyniel, Sigman, et al., 2015): akin two sides of the same coin, the former describes the likelihood a chosen action will produce a desired outcome and the latter describes the complementary probability it will produce any outcome, but the desired one. Here, we aimed at investigating the neural dynamics underlying the update of these estimated probabilities or beliefs, with or without explicit value-based outcomes. To this end we developed two multi-option decision-making tasks (Fig. 1) to allow non-binary decision-making (Churchland & Ditterich, 2012; Churchland, Kiani, & Shadlen, 2008; Tajima, Drugowitsch, Patel, & Pouget, 2019). We used a multi-option design instead of the traditional two-option setup to avoid the possibility of participants using any exclusion/confirmation criterion as a strategy for decision-making. The tasks also employed three categorical types of data and identical volatility rate (i.e. average number of trial before reversal), therefore conflating noise and surprise (McGuire, et al., 2014; Nassar, et al., 2016; Nassar, McGuire, Ritz, & Kable, 2019) and providing participants with discrete evidence for the belief update. This design fostered a constant rate of belief updating (S. Lee, Gold, J. I., & Kable, J. W., 2020; Soltani & Izquierdo, 2019), allowing for nuanced transitions between trials characterized by high confidence (i.e. one choice selection associated with the desired outcome with high probability) and those characterized by high uncertainty (i.e. chance level to produce the desired outcome, distributed across all available choices). Two similar Bayesian learner models were employed to estimate subject-specific, trial-by-trial, belief updating and associated choice-related confidence/uncertainty, replicating the actual behavior recorded in healthy volunteers (N=28) being scanned with functional magnetic resonance imaging (fMRI).

**Figure 1.**
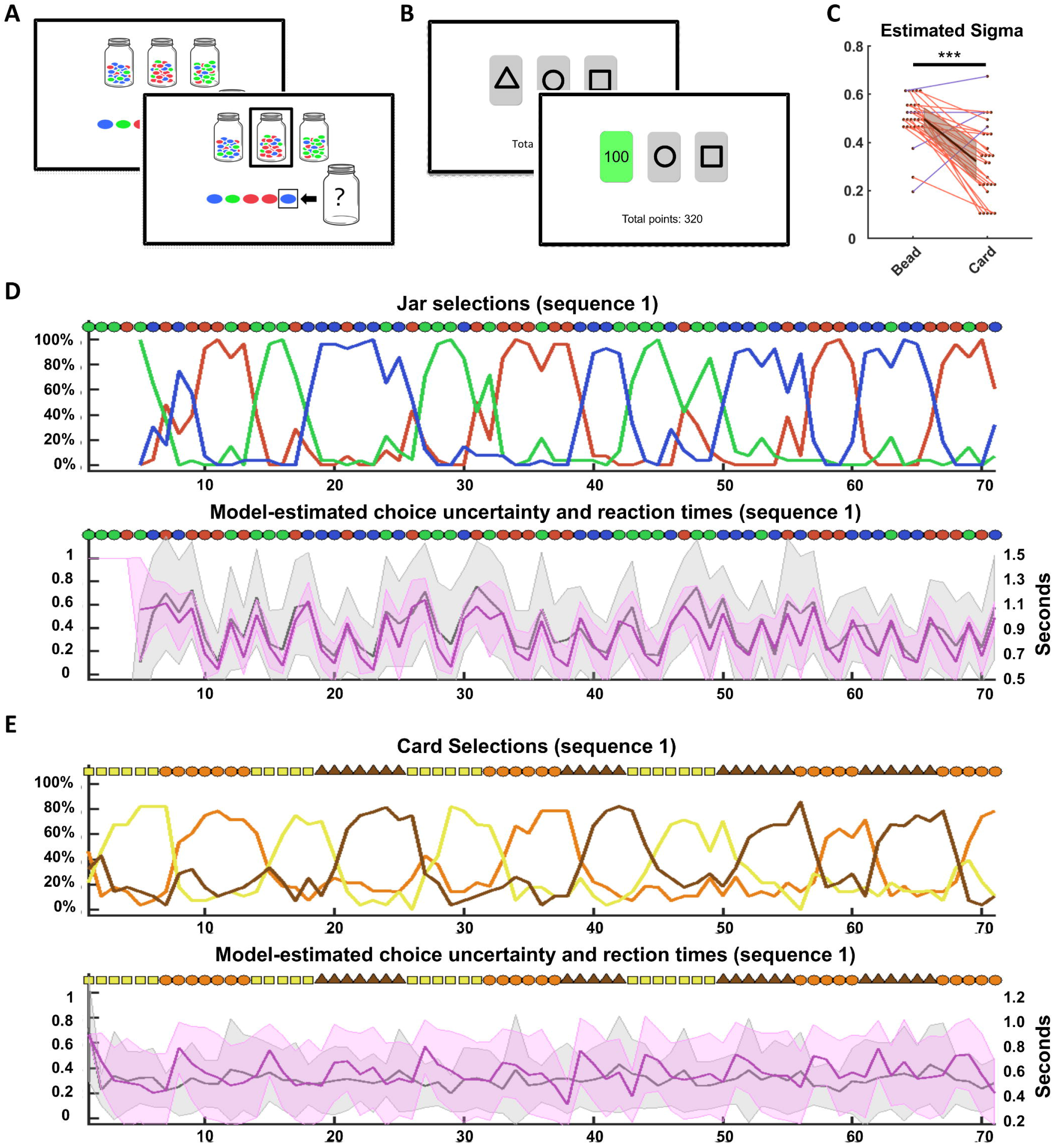
Experimental paradigm, choice behavior, and model estimations for trial-by-trial confidence and uncertainty. (**A**) Bead task. Participants chose from which of three jars, containing predominantly red, blue or green beads (80%-10%-10% ratio), the latest bead was extracted from. The latest five extracted beads were always displayed on screen, and no feedback was provided after each choice selection. (**B**) Card task. Participants chose one card among three cards characterized by different geometric figures. Each card yielded 100, 10 or 0 points (immediate value-based outcome) with a probability distribution of 80%-10%-10%. The extraction jar and the card-value associations were changed every 5, 6 or 7 trials, in three pre-established pseudo-random sequences that were used for all participants, in a counterbalanced order. (**C**) Subject-specific values of the coefficient σ (grouped in intervals of .03 in the scatterplot), used in the Bayesian models to estimate confidence and uncertainty in each participant on a trial-by-trial basis. Different σ values determine different paces of belief update: for instance, the higher the value, the more confirming evidence is required to increase confidence and decrease uncertainty. The color of each extracted new bead and the feedback provided after each card selection was used as discrete new evidence to update the estimated probability distribution or priors of each participant. (**D-E**) Choice selections expressed by the participants (upper row, as percentages) and model-estimated choice-uncertainty (magenta, lower row, expressed as mean and standard deviation). In grey, superimposed on the estimated choice uncertainty, mean and standard deviation of the reaction times, per task. ***: p<.001

Previous investigations have revealed a number of brain regions involved in the computations of belief updating and the associated decision confidence and uncertainty (Meyniel & Dehaene, 2017; Morriss, Gell, & van Reekum, 2018; Pouget, et al., 2016). For instance, the anterior insular (AIC) (Gu, Wang, et al., 2015; Palminteri, et al., 2012; Seymour, et al., 2004) and dorsal anterior cingulate (dACC) (Behrens, et al., 2007; Botvinick, 2007; Rushworth & Behrens, 2008) cortices have been shown to monitor prediction errors and conflicts in general; the dorsolateral prefrontal cortex (dlPFC) (Glascher, Daw, Dayan, & O’Doherty, 2010; S. W. Lee, Shimojo, & O’Doherty, 2014; McGuire, et al., 2014) has been reported to process reward-independent state prediction errors. Furthermore, the medial prefrontal cortex (mPFC) (Bang & Fleming, 2018; Matsumoto & Tanaka, 2004; Yoshida & Ishii, 2006) has been shown to be responsible to process self-monitoring, action evaluation and choice confidence in goal-oriented behavior. At the subcortical level, the anterior hippocampus (aHip) (Harrison, et al., 2006; Rigoli, Michely, Friston, & Dolan, 2019; Strange, et al., 2005) has been implicated in monitoring expectations, predictability and entropy reduction in changing environments, whereas the amygdala (Amg) (Bechara, Damasio, Damasio, & Lee, 1999; Dolan, 2007; Jung, Lee, Lerman, & Kable, 2018) has been associated with value representation and risk estimation in stochastic environments. Consistent with this literature, we found two neurocircuitries supporting belief updating across both tasks: the first network, encompassing mPFC, hippocampus and amygdala, became more active as decision confidence increased; the second, which included AIC, dACC and dlPFC, become activated as decision uncertainty built up. We hypothesized network dynamics would be affected by the availability of explicit outcomes, therefore we used dynamic causal modeling (DCM) (K. J. Friston, Harrison, & Penny, 2003; Stephan, et al., 2010; Zeidman, et al., 2019) to quantify the directed influence or effective connectivity within each of the two active neurocircuitries.

## Materials and Methods

### Participants

We recruited 28 healthy volunteers (16 females), age 24.8 ± 7.0. Participants taking any medication, or with a history of mental disorder or drug abuse were excluded from the study. One subject was excluded from all fMRI analysis involving the card task, due to excessive movement. The study was approved by the Institutional Review Board at the University of Texas, Dallas and University of Texas Southwestern Medical Center. Informed written consent was obtained from all subjects and all participants were informed that they could withdraw from the study at any point.

### Experimental design: 3-options continuous choice tasks

In a first task (**Fig. 1A**), termed *bead task* (modified from: Huq, Garety, & Hemsley, 1988), a new visual stimulus (a red, blue or green *bead*) was presented at each trial, adding a congruent (e.g. a red bead after another red one) or incongruent (e.g. a blue bead following a green one) color stimulus in a sequence of beads. Participants had 2 seconds to decide from which of three jars displayed on the monitor the bead had been drawn. The jars were illustrated on screen as containing beads of three colors in a ratio of 80%-10%-10%. After each button press, the selected jar would be highlighted with a black rectangle, for the remaining time on the clock allowed for the choice selection, plus 0.5 seconds. The last five extracted beads were always present on screen (**Fig. 1A**), and the participants were allowed to make the first choice selection starting from the 5^th^ bead extraction. Between trials, a grey square would appear to conceal the new extracted bead for a variable time of 2 to 3.5 seconds. Participants had access to immediate value-based outcomes during a training session, which would show that each correct guess would result in accumulating 100 points. However, no outcome was provided during the MRI task and the participants were made aware that the accumulated points would be disclosed only at the end of the task.

In the second task, termed *card task* (**Fig. 1B**) (modified from: J. O’Doherty, Kringelbach, Rolls, Hornak, & Andrews, 2001), the participants had 2 seconds to select among three cards presented on the screen and characterized by three geometric figures (randomly selected among triangle, square, circle, star and diamond). Each card was assigned a different predominant value among the three possible outcomes of 100, 10 and 0 and after selecting a card, the screen would display a stochastic outcome, highlighted in green (variable duration: 1-2.5 seconds), with assigned probability of 80%, for the predominant value, and 10% for each of the remaining outcomes. A second message on screen also signaled the amount won on white screen (fixed interval: 0.5 seconds), followed by a fixation cross (variable interval: 1-2.5 seconds), which would precede a new trial. The total amount of points accumulated, per block, was always displayed in the lower part of the screen.

Finally, the participants were instructed that the computer would randomly change the jar from which it extracted beads, or the card-value associations. The pace of these pseudo-random changes was determined in an interval of 5 to 7 trials for both tasks and it was independent of the performance of the participants (**Fig. 1D,E**). For both tasks, participants were compensated with $1 for every 500 points, selecting the points accumulated during one random block per task. Both tasks consisted in 3 blocks of 71 trials each, so the participants were told that the maximum amount of bonus they could gain consisted in about $15 dollars from each task. Three identical sequences -one per block-were used for all subjects, for both the bead colors and the card-value associations. Task order and block order were counterbalanced across subjects.

### Bayesian learner model

We used two similar computational models to estimate: 1) in the bead task, the trial-by trial subjective probability assigned to each jar, as the source of extraction of the latest bead visible on screen; 2) in the card task, the trial-by trial subjective probability that each card would be associated with the highest chances to receive a 100 points reward. We then used the model-estimated probability assigned to the selected jar or card, per trial, to determine subject- and trial-specific choice-confidence (*c*), with the complementary uncertainty (*1-c*).

To update the estimated subjective probabilities the model relied on a Markov chain: at the beginning of each trial *t*, the estimates defined at time *t-1* were incrementally updated into the new probabilities, depending on the latest available evidence and the subject-specific assumptions about the likelihood of events in the environment. For instance, if one assumed the environment were characterized by low stochasticity (i.e. vast majority of beads in the jars are of a single color and most feedbacks after a card selection match the assigned value), she would quickly adapt to any change of bead color or card selection outcome, i.e. the behavior relies largely on the latest presented evidence. Conversely, an environment assumed to be characterized by high stochasticity resulted in a slow adaptation of estimated probabilities, as conflicting evidence had to accumulate to trigger a change in choice selection, i.e. the behavior relies mostly on multiple previous estimates. These distributions of events are defined in Bayesian terms as likelihood, or the probability of an event to occur, assuming an hypothesis is true: e.g. p(bead_blue_ | ExtractionJar_blue_) or p(reward_100_ | Card_100_). Participants were not aware of the exact distribution of colored beads in the jars or outcome stochasticity associated with card selections. Therefore we considered each subject would rely on their own assumptions about these likelihoods, which would be kept constant through the task, as the task instructions explicitly mentioned the probability distributions would not vary depending on performance or other measures. The model assumed that the participants relied on a normal distribution to describe the likelihood of the events in each task (i.e. distribution of beads, across jars, or stochastic values, across cards):

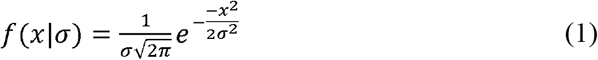

Where σ represents the standard deviation of the distribution and x=[-1 0 1] is normalized to determine the likelihood assigned to all the three possible events associated with each hypothesis (e.g. p([bead_red_ bead_blue_ bead_green_] | ExtractionJar_blue_) or p([10_points_ 100_points_ 0_points_] | Card_100_)), where the most likely event was assigned the value of the mean of the distribution. For instance, a σ value of 0.5 resulted in a likelihood of 78.7% for the most likely event and 10.65%, equally distributed between the two remaining events, whereas a σ value of 0.1 resulted in a likelihood of ≈100% for the most likely event and ≈0% for the remaining events. Thus, subject-specific σ values resulted in different paces of belief updates, in the face of new trial-by-trial data (newly extracted bead or selection reward): the higher the value of σ, the higher the stochasticity assumed by a subject in the environment, and therefore the slower the pace of belief update. This update process can be summarized as:

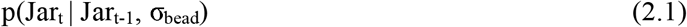

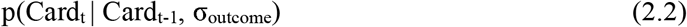

For both tasks, the update process followed a standard Bayes rule:

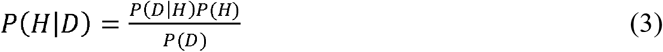

In the context of the two tasks, H (the hypothesis) represents the jars for the bead task and the cards for the card task. Similarly, D (the data) represents the color of the latest extracted bead, for the bead task, and the value of the latest outcome, for the card task. *P(H)* indicates the probability (carried from a previous trial) assigned to each jar or each card to be the extraction jar or the card assigned 100 points. *P*(*D*) represents the total probability of the observed data to occur, given the subject-specific assumptions about the distribution of events in the task environment and the carried probabilities assigned for P(*H*). As a result, for the Bead task, a new colored bead observation resulted in increasing the probability associated with the matched colored jar and decreasing the probability associated with the remaining ones. Similarly, for the card task, an outcome equivalent to 100 would increase the probability to find the same outcome again by re-selecting the card that delivered the 100 outcome. Conversely, outcomes of 10 or 0 would decrease the probability to find the maximum reward by repeating the same choice, increasing instead the probability associated with both non-selected cards. Finally, the updated three dimensional vectors of probabilities in each task were transformed – after the calculation of the error required for parameter regression-to avoid any selection to reach 0% probability, which would have hindered future incremental updates, as follows: max[.05, Jar_t_] and max[0.05, Card_t_].

We used a Monte Carlo method (i.e. random search within the space of parameters) to estimate the value of the parameter σ, per each task, that would better match real choice selections expressed by each of the 28 participants. For the regression, we coupled the trial-by-trial model-estimated distribution of probabilities across the three available choices with the actual choice selection of each participant. This value (*c*) was used to generate an error score per trial, which was computed as |log(*c*)|. The regression method tested 10^3^ randomly generated values for the parameter σ, in a [0 1] interval, searching for the value that minimized the total error across all sequences, per task (Fig. 1c).

### Alternative computational models

The performance of the Bayesian learner model was compared with three more computational constructs, all based on a similar Markovian principle of trial-by-trial update.

In a first model, we employed a simple heuristic that would increment or decrement the probability associated with a jar or a card by a subject-specific constant *H*:

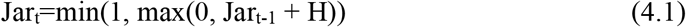

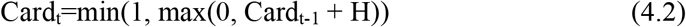

where *H* is a vector 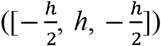, so that in the bead task the Jar of the same color of the latest extracted bead would increase its estimated probability by a value of *h*, whereas the remaining two Jars would decrease their probability by 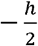 Similary in the card task, a 100 point outcome would trigger an increase of probability for the selected card, decreasing the probabilities assigned to the non-selected cards, whereas the opposite process was employed for 10 points and 0 points outcomes. The vectors of estimated probability were normalized at each trial and the value of *h* was regressed for each subject (maximum log likelihood) to find the value that would reduce the model-estimated error.

In a second model, we employed a Bayesian learner similar to the one already described, with the key difference of relying on a dynamic pace of update, rather than a fixed one (cf. Behrens, et al., 2007). To this end, the value of the parameter σ (equation 2) was itself estimated trial by trial as a function of the estimated volatility of the environment:

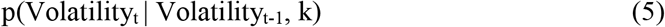

where the value of k controlled the pace of change of volatility and was estimated in each subject to match the actual behavior of the participants. Due to this change in the Bayesian learner model, the value of σ increased under condition of low volatility (i.e. a sequence of beads of the same color, or a sequence of 100 point reward) and decreased otherwise, thus dynamically adjusting the pace of behavioral adaptation.

In a third model we relied on a reinforcement learning approach, employing the learning rule:

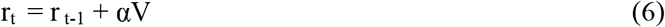

where *r* represents the expected reward or outcome, per trial, α is the learning rate, estimated in each subject, and V is the prediction error, computed with a standard Rescorla-Wagner learning rule. In the Bead task, where no explicit reward is provided, we assumed that a bead at trial *t* of the same color of the chosen jar at trial t-1 would be considered by the participant as an outcome of 100 (i.e. confirming the previous choice was correct). Conversely, an inconsistency between the present bead color and previous choice of jar color would be encoded as an outcome of 0. In both tasks, choice selection was then determined using a linear transformation (cf. Behrens, et al., 2007):

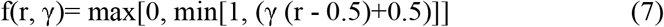

The parameter γ, which was also estimated in each subject, represents whether the participant was risk averse (>1) or risk prone (<1) in their choice selections.

In conclusion, we compared the predictions of the models in terms of accuracy (i.e. mean estimated probability associated with trial-by-trial choice selection) and BIC score. Limited to the model comparison, to allow for comparable BIC scores, we used a softmax transformation of the trial-by-trial estimated probability (τ=.1 for all models) and relied on the maximum log-likelihood (controlled by the Matlab function *fmincon*) for the parameter regression, across models. The Bayesian learner used for the fMRI and DCM analysis outperformed all the other tested models in both tasks (table 1).

**Table1:**
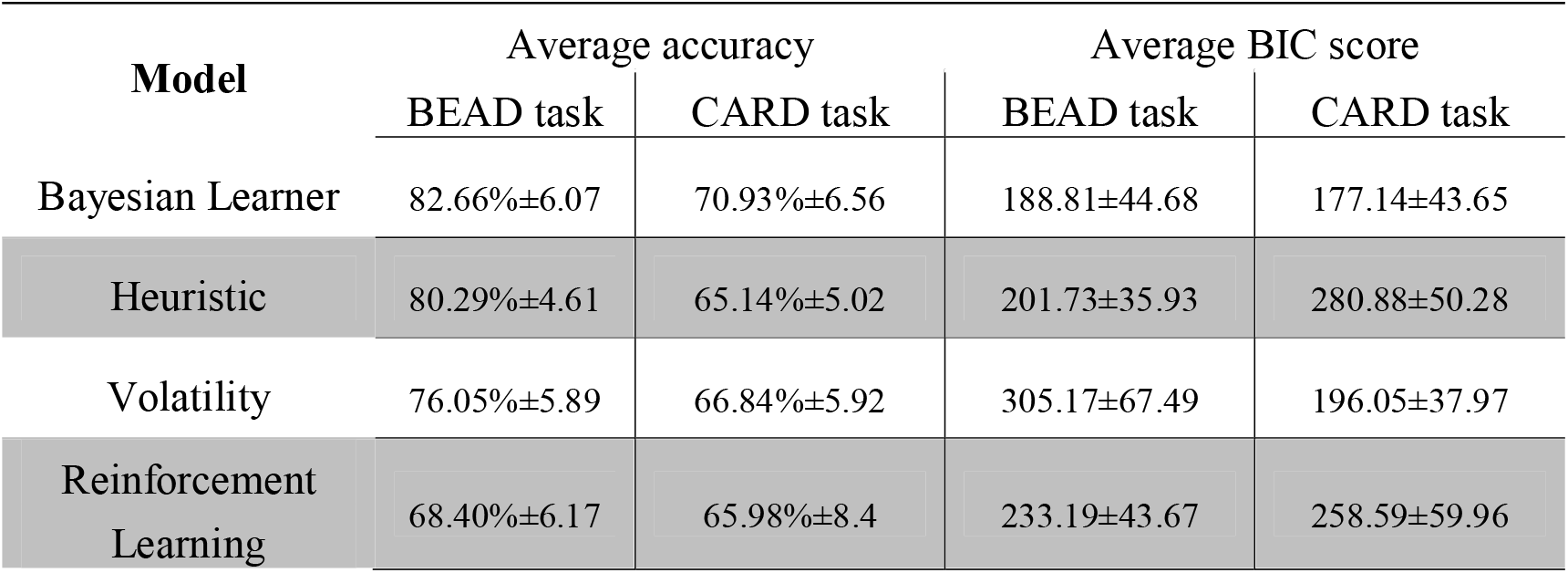
Model comparison

### fMRI data acquisition and preprocessing

Functional MRI data were acquired using a Philips 3-Tesla MR scanner at the Advanced Imaging Research Center at University of Texas Southwestern Medical Center. The anatomical scan sequence (multi-echo MPRAGE) was carried out with a resolution of 1 mm, Multi Parametric Maps. Functional images (EPI) were acquired with a resolution of 3.4 x 3.4 x 4mm, repetition time of 2000 milliseconds, echo time of 25 milliseconds, 38 axial slices, flip angle=90°, and a field of view of 240 mm. We used standard Statistical Parametric Mapping algorithms (SPM12, Wellcome Department of Imaging Neuroscience; www.fil.ion.ucl.ac.uk/spm/) for data preprocessing, including motion realignment to the first volume, coregistration to the participant’s anatomical scan, MNI normalization, and spatial smoothing, using an isotropic 8-mm full-width at half-maximum (FWHM) Gaussian kernel.

### Model-based fMRI

We considered choice confidence and uncertainty encompass several steps in a process, each responsible for part of the probability estimation in a choice selection (Ma & Jazayeri, 2014; Meyniel, Sigman, et al., 2015), from sensory estimation to motor execution (Orban & Wolpert, 2011; Wolpert & Landy, 2012), or outcome evaluation (Bach, Hulme, Penny, & Dolan, 2011; Meyniel, Schlunegger, et al., 2015), when at all available. Thus, for the GLM analysis, we used default SPM12 functions to observe BOLD signals associated with the onsets of choice selections (i.e. button presses). This is consistent with our computational definition of confidence and uncertainty (Meyniel, Sigman, et al., 2015), which was meant to investigate both these phenomena in association with a choice, i.e. the estimated probability a selected action would deliver a desired result. We convolved a canonical hemodynamic function (HRF), which is a synthetic hemodynamic response function composed of two gamma functions (K. J. Friston, et al., 1998; K.J. Friston, et al., 1994) in SPM, with task regressors related to all onset of choice selections. In each task, we used two separate GLM analyses to identify the relationship between the -parametrically modulated-task events and the hemodynamic response. For these analyses, we used the RTs as trial-by-trial covariates and either choice confidence (c) or choice uncertainty (1-c) as parametric modulators. The three blocks were concatenated in both task. Whole brain activations were determined using a threshold of *P*<.05 corrected for familywise errors (*P_FWE_*<.05; **Fig. 2A,B**).

**Figure 2.**
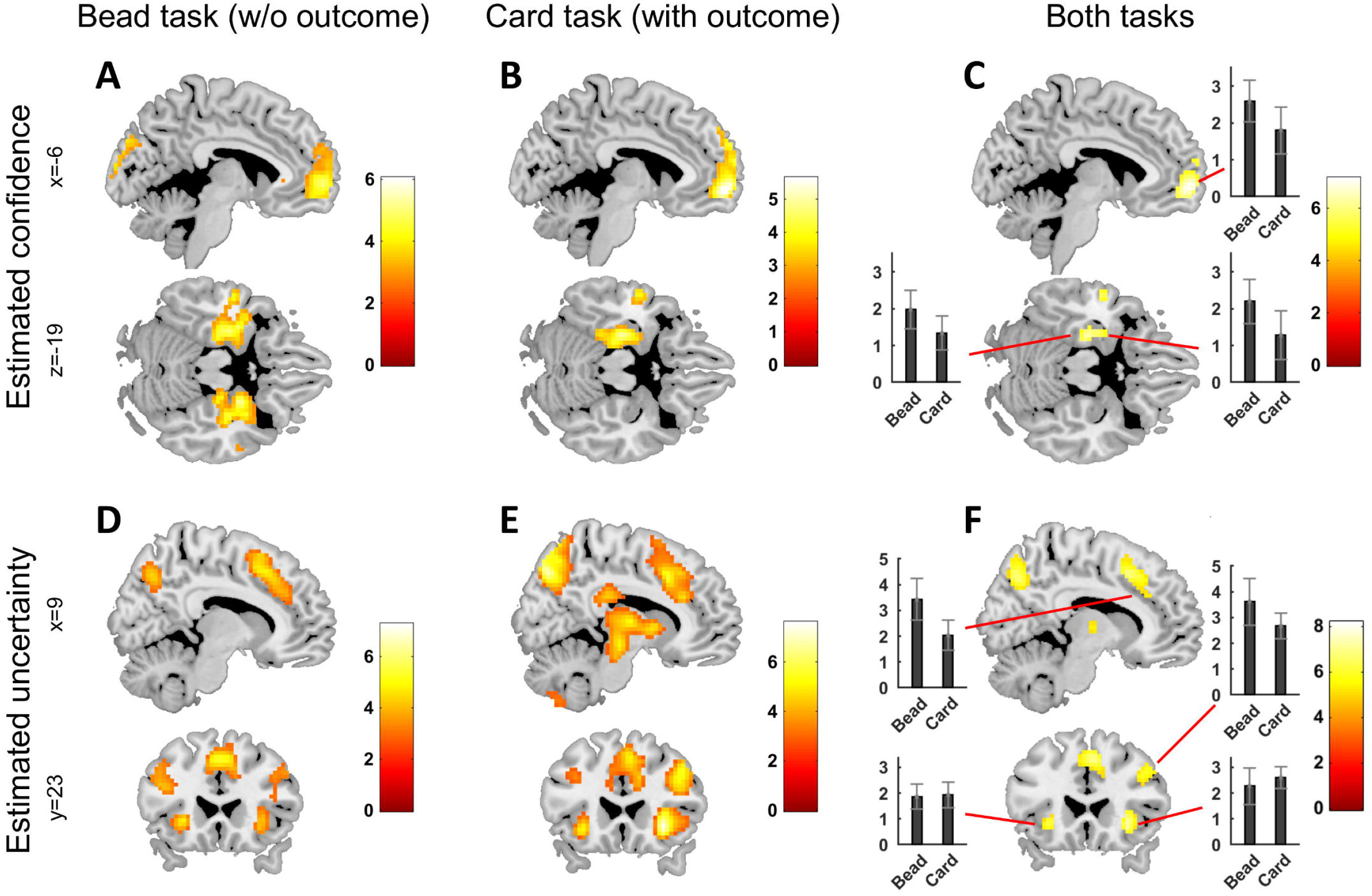
Model-based BOLD responses encoding Bayesian confidence and uncertainty. (A-C) fMRI BOLD response recorded using the model-estimated signal of confidence (*c*) as parametric modulator and reaction times as covariates, in the bead task (A; *P_uncorrected_*<0.005, k=50), the card task (B; *P_uncorrected_*<0.005, k=50) and considering both tasks together (C; *P_FWE_*<0.05). (D-F) fMRI BOLD response recorded using the complementary model-estimated signal of inverse confidence or uncertainty (*1-c*) as parametric modulator and reaction times as covariates, in the bead task (D; *P_uncorrected_*<0.005, k=50), the card task (E; *P_uncorrected_*<0.005, k=50) and considering both tasks together (F; *P_FWE_*<0.05). (C, F) Bar plots and standard error of the mean illustrating *β* values extracted from bead and card task across multiple regions of interests.

### Dynamic causal modelling (DCM)

For the DCM estimation, we relied on SPM12 default functions. Our model-based GLM revealed two distinct networks subserving uncertainty (AIC, dACC, dlPFC; **Fig. 2A,C,E**) and confidence (aHip, Amg, mPFC; **Fig. 2B,D,F**). We extracted fMRI time series from individual ROIs, using their principal eigenvariates: we relied on anatomical maps manually generated for the Amg and the aHip, and we used spherical ROIs (8mm radius) for all cortical regions. These were centered on task-specific, group-averaged, peaks activity, at the following coordinates: AIC (bead: [-27, 23, −1] and [33, 23, 2]; card:[−30, 20, −7] and [30, 23, −4]), dACC (bead: [−6, 29, 41] and [6, 29, 41]; card:[-9, 29, 23] and [9, 29, 26]), dlPFC (bead: [−42, 23, 26] and [42, 29, 29]; card:[−36, 26, 32] and [36, 26, 29]), mPFC (bead: [−9, 56, −1] and [9, 56, −1]; card: [−9, 56, −4]). We also included the visual cortex as a network input region (cf. K. J. Friston, et al., 2003; Stephan, et al., 2007; Stephan, et al., 2008), with ROIs centered on the following coordinates: bead: [12 −88 32] and [−12 −94 −7]; card: [21 −91 11] and [−21 −91 11]. The time series extracted in the visual cortex ROIs were derived from a baseline BOLD activity, before the use of covariates and parametric modulators. With the exception of the response to the signal of confidence recorded in the card task, which was limited to the ROIs in the left hemisphere, we found significant increase in BOLD response bilaterally, for all ROIs. Thus, to allow comparison across tasks, we focused the DCM analysis for the signal of confidence on the left hemisphere and for the signal of uncertainty on both hemispheres. The latter was used also as cross validation of the results within tasks.

For the network architectures, we restricted the DCM analysis to those architectures that would better inform about changes in directional relationships among the active ROIs. This was not meant to try to exhaust all possible model structures, but rather to aim at a good balance between accuracy and complexity, thus affording a sufficient generalizability (Pitt & Myung, 2002; Stephan, et al., 2010). Therefore, we compared neural architectures that fulfilled two criteria: 1) they presented different targets for the modulatory signals; and 2) they were computationally comparable in terms of the number of free parameters (i.e. comparable number of node-to-node and modulatory connections (cf. Yu, et al., 2020)). These two requirements led to develop eight models, which were all characterized by a fully connected *A* matrix and a *C* Matrix directing the screen-based sensory input towards the Visual cortex node (Gu, Eilam-Stock, et al., 2015; e.g. see: Gu, Wang, et al., 2015). Then, we established the targets of the modulatory inputs (the *B matrix*), represented by the signal of estimated confidence (*c*) or uncertainty (*1-c*). We assumed that these modulatory inputs would affect the connectivity between visual cortex and the main three active ROIs, bilaterally, for all models, but they would target the connectivity between the three main ROIs in one direction only. Thus, the eight models did not differ in the overall number of targets of modulatory inputs (6 fixed and 3 variable), but in the combination of selected targets (2 possible directions between 3 pairs of nodes, **Fig. 3A-B**).

**Figure 3.**
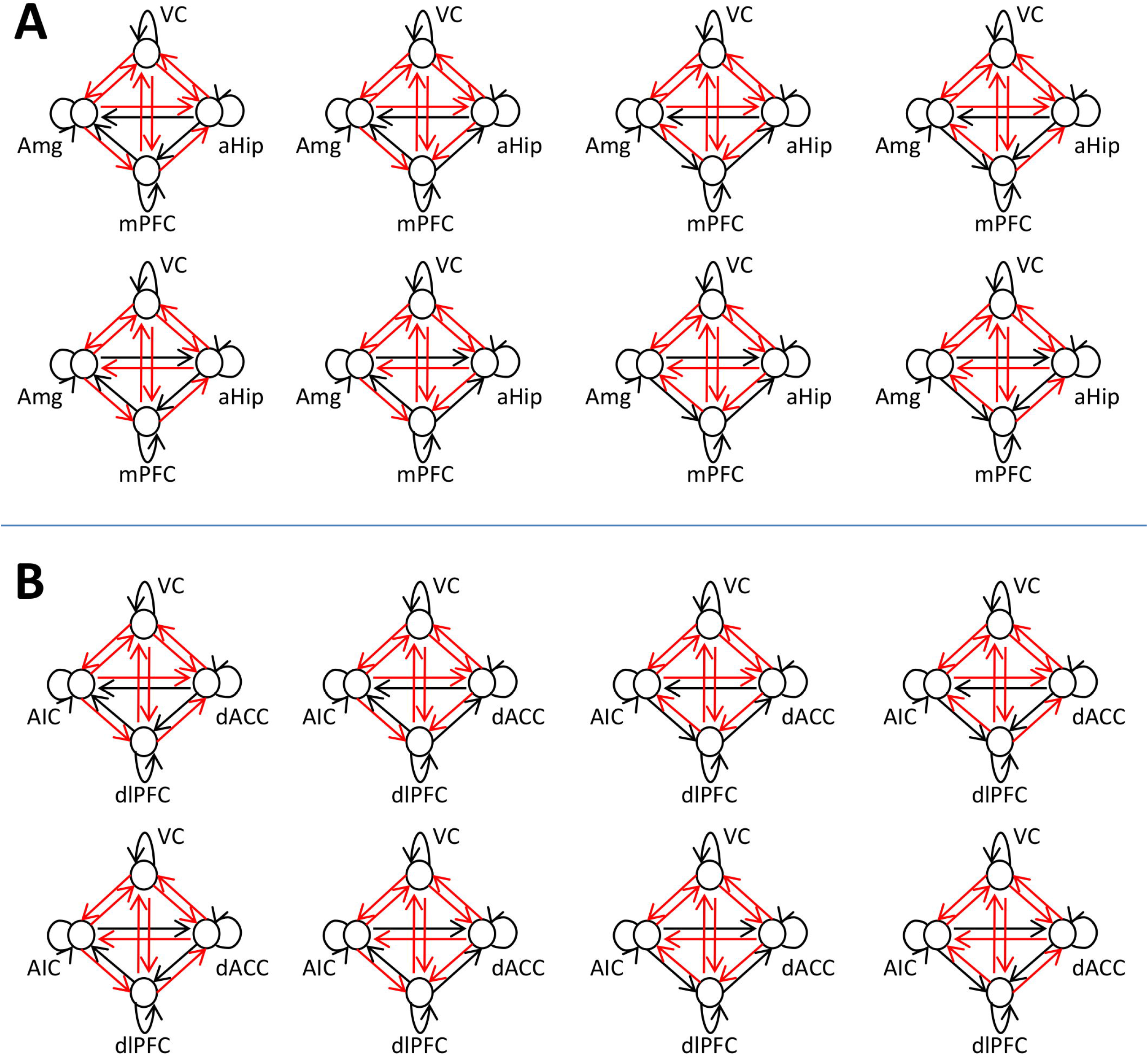
Neural network architectures tested in the family-wise DCM. These neural architectures illustrate the eight models used to test the effective connectivity associated with the signal of confidence (A) or its complementary probability, uncertainty (B). The same model structure have been tested in both tasks to ease a comparison of the results. The general architecture of connectivity (*A matrix*) remains unchanged and it is represented by all the arrows in the network illustrations. Conversely, the targets of the modulatory inputs (*B matrix*), marked only by the red arrows, are different in each model. This differentiation results in the generation of the 8 models, then divided into competing families of 4, to allow family-wise comparisons for each pair of nodes in the ROI triplets aHip-Amg-mPFC or AIC-dACC-dlPFC. For instance, to determine whether the directed modulated connectivity from Amg to aHip better explains the extracted data, in comparison with the aHip-to-Amg connectivity, the top row of four models in panel A are considered in a single family against the lower row of four models in the same panel. (aHip: anterior hippocampus; AIC: anterior insular cortex; Amg: amygdala; dACC: dorsal anterior cingular cortex; dlPFC: dorsolateral prefrontal cortex; mPFC: medial prefrontal cortex; VC: visual cortex).

We used a Bayesian family-wise random effect approach to estimate the directed connectivity between each pair of nodes within the two network triplets, grouping the eight models each time into two families of four models, each family characterized by one common modulated directed connectivity. This method allowed comparing, for instance, the four models with an AIC-to-dACC modulated connectivity (with varying connectivity between AIC and dlPFC and between dACC and dlPFC, **Fig. 3A**) vs the four models characterized by an dACC-to-AIC modulated connectivity (and varying connectivity for the remaining couples of nodes). The method allowed to assess the likelihood a modulatory signal would affect the information flow in a specific direction, within a single pair of nodes, independent of the presence of a single winning model that would have emerged in a single model comparison. Finally, we tested our results independently across tasks and, limited to the uncertainty signal, across hemispheres within each task.

## Results

The Monte Carlo regression resulted in subject-specific values determining each participant’s own pace of belief update (σ=.497±.095, σ=.321±.156, respectively for the bead and card task, Fig. 1C). Within-subject comparison revealed the bead task was characterized by significantly higher σ values (i.e. slower pace of belief update) in comparison with the card task (d(27)= 5.39, p<.0001). These values resulted in optimized Bayesian models that provided a mean behavioral prediction accuracy of 82.66%±6.07% for the bead task and 70.93%±6.56% for the card task (chance≈33%).

A correlation analysis revealed that the estimated confidence (*c*) was negatively correlated with the reaction times (RTs) in the bead task for 27 participants out of 28 (mean *r* = −.44 ± .15, range: [−0.2, −0.7], *P*<=.004, for all significant correlations), whereas in the card task only five participants showed a significant (*P*<.05) correlation (cf. **Fig. 1D and E**). Next, we included RTs values as covariates and we examined the neural activations associated with either the estimated confidence (*c*) or uncertainty (1-*c*) as parametric modulators in separate GLMs (**Fig. 1D,E**). Consistent with previous literature (Meyniel & Dehaene, 2017; Morriss, et al., 2018; Payzan-LeNestour, et al., 2013; Pouget, et al., 2016), we found that confidence was encoded in the mPFC, left aHip and left Amg, across both tasks (**Fig. 2A-C**, *P_FWE_*<.05). The inverse of confidence, or uncertainty, was encoded in bilateral AIC, dACC, and right dlPFC, across tasks [**Fig. 2D-F**; *P*<.05, family-wise error (FWE) corrected]. A within subject comparison of *β* values extracted from these nine ROIs across tasks confirmed the strong similarities of BOLD neural responses, independent of the presence of explicit value-based outcomes. Only the activity in the left dACC revealed a difference between tasks (d(26)=2.48; p=.020; bead>card) which would not survive after correcting for multiple comparisons (α<.0056). Finally, a whole brain contrast between the two tasks revealed increased activity (peak: [27 47 20]) in the right lateral frontopolar cortex (Koechlin & Hyafil, 2007; Mansouri, Koechlin, Rosa, & Buckley, 2017), in association with uncertainty processing, when comparing Card vs Bead task BOLD activity (cluster size>50, *P*<.005, uncorrected), but this result would not survive FWE correction.

Next, we used DCM to investigate the directional dependencies among confidence- and uncertainty-encoding regions. Family-wise model comparison was used to test pair-wise directed connectivity among Amg, aHip and mPFC, in the left hemisphere across tasks, or AIC, dACC and dlPFC, in both hemispheres across tasks, grouping the eight tested models (**Fig. 3**) into two competing families of four models, depending on the tested connectivity. This analysis revealed that the directed influences associated with the signal of confidence were found to change depending on value-based outcome availability. In the bead task, when immediate value-based outcome was absent, confidence primarily modulated the connections from the aHip to other regions (exceedance probability, left hemisphere: aHip-to-mPFC: 94% and aHip-to-Amg: 97%; **Fig. 4A**), with no clear directionality for the remaining pair-wise analysis (exceedance probability, left hemisphere: mPFC-to-Amg: 54%). In contrast, in the card task, which provided immediate value-based feedback to each choice, the signal of confidence primarily modulated mPFC-to-aHip, mPFC-to-Amg, and Amg-to-aHip connectivity (exceedance probability, left hemisphere: 95% and 81%, and 82%, respectively; **Fig. 4B**).

**Figure 4.**
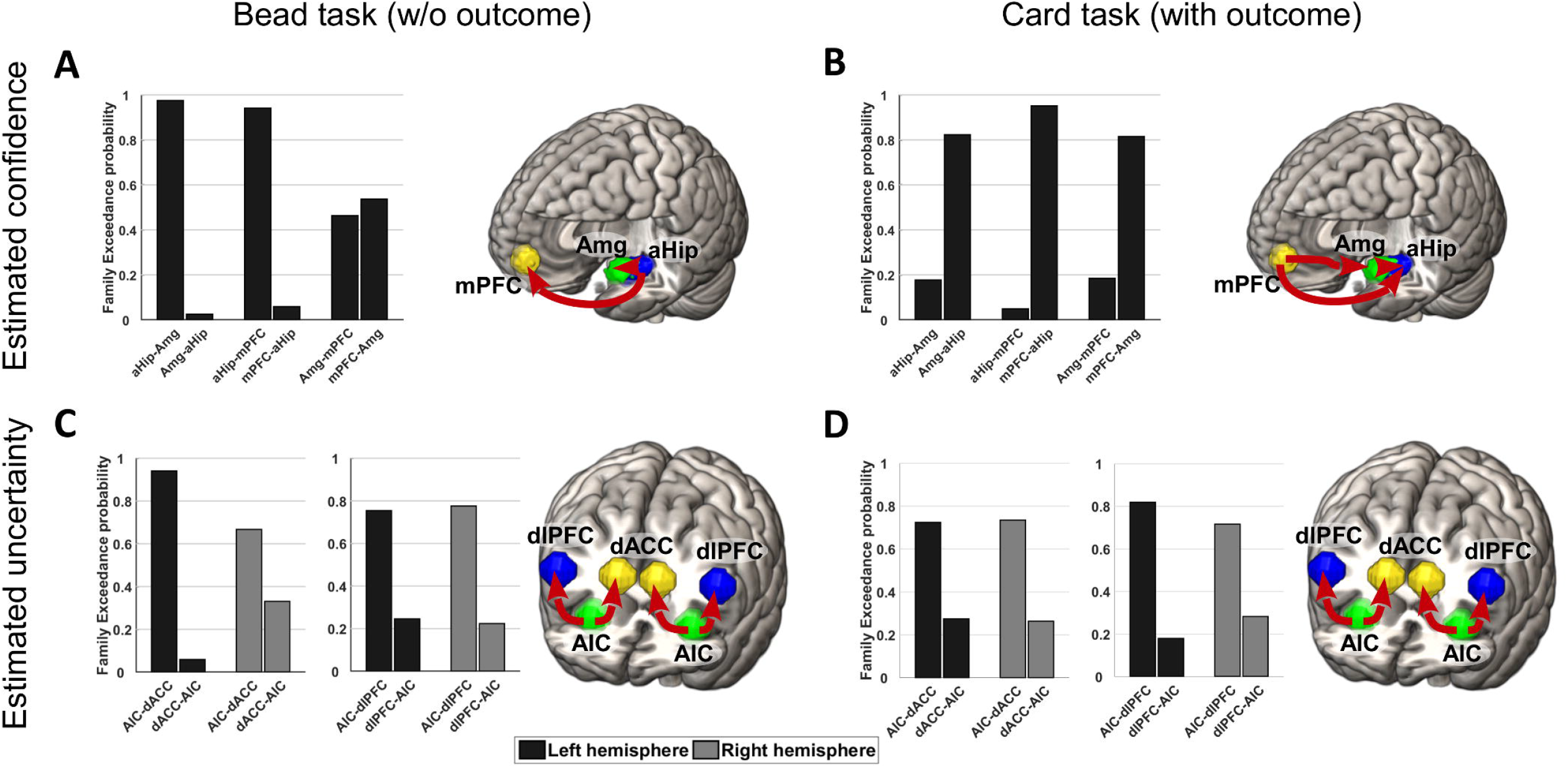
DCM results using family-wise comparison. Histograms report the family-wise Exceedance probability for each tested pair-wise connection in each task. The network diagrams illustrate a summary of the pair-wise analysis, with thick red arrow used to highlight the estimated direction of effective connectivity. Missing connectivity between nodes represent those pair-wise analysis for which DCM did not provide a conclusive result. In the bead task the aHip drives the confidence-building network (**A**), whereas in the card task mPFC and Amg drive the activity of the aHip in association with increased signal of confidence (B). The uncertainty-building network remains unchanged in the comparison across tasks and it is driven by the activity of the AIC both in the bead task (C) and in the card task (D). (aHip: anterior hippocampus; AIC: anterior insular cortex; Amg: amygdala; dACC: dorsal anterior cingular cortex; dlPFC: dorsolateral prefrontal cortex; mPFC: medial prefrontal cortex)

Interestingly, our DCM analysis also suggested that the neural dynamics within the uncertainty network did not vary as a function of outcome availability. Specifically, the signal of *uncertainty* modulated AIC-to-dACC [exceedance probability for bead task: 94% (left hemisphere), 67% (right hemisphere), **Fig. 4C**; card task: 72% (left),74% (right), **Fig. 4D**] and AIC-to-dlPFC connectivity [exceedance probability for bead task: 75% (left),78% (right), **Fig. 4C**; card task: 82% (left),72% (right), **Fig. 4D**]. Finally, the DCM analysis did not indicate a clear direction in the dlPFC-dACC effective connectivity, as results did not converge across hemispheres and tasks [exceedance probability for bead task: 69% (left hemisphere), 47% (right hemisphere); card task: 63% (left), 59% (right)]. The converging results allowed to run ttest comparisons for the weight of modulatory connectivity estimated in the two tasks, limited to the AIC-to-dACC and AIC-to-dlPFC connectivity, across hemispheres (Stephan, et al., 2010). These comparisons did not reveal any significant difference, after correction for multiple comparisons (Bonferroni correction, α=.0125).

## Discussion

Humans live in constantly changing environments that are often opaque, as explicit (e.g. hedonic or value-based) outcomes following a choice behavior is not always available. Our main findings revealed two important aspects of belief updating in uncertain environments with or without immediate outcomes. First, we identified two opposing networks responding to either decision confidence (amygdala, anterior hippocampus, mPFC) or decision uncertainty estimations (AIC, dACC, and dlPFC), regardless of the presence of immediate value-based outcomes. Second, DCM analysis indicated that the network dynamics changed as a function of feedback availability only for the confidence-encoding network. Taken together, these findings revealed important changes across different Marrian levels of analysis (Marr & Poggio, 1976), as the presence of immediate value-based outcomes impacted the neural implementation of belief updating in its confidence-, but not uncertainty-building, component, despite the identical computational definitions for the two measures across condition of feedback access.

Existing literature on decision-making primarily focuses on choice made in environments with explicit feedback consisting in value-based outcomes, assuming previously experienced outcomes following chosen actions are needed to generate subjective values and to guide future choices (Berridge & Kringelbach, 2015; J. P. O’Doherty, Cockburn, & Pauli, 2017; Rangel, et al., 2008). This approach has been highly successful in accounting for different forms of conditioning, habitual and goal directed behavior and in uncovering their underlying neural substrates (Balleine, Delgado, & Hikosaka, 2007; Balleine & O’Doherty, 2010; Dezfouli, Lingawi, & Balleine, 2014). The algorithmic formalization offered by the reinforcement-learning framework (Sutton & Barto, 1998) further expanded the domain of decision-making investigations in accessible environments, indicating reward prediction-error are encoded by dopamine signals (Schultz, 2002; Schultz, et al., 1997), and driving model-based analysis of neural activity (Daw, Gershman, Seymour, Dayan, & Dolan, 2011; Daw, Niv, & Dayan, 2005; Dolan & Dayan, 2013; S. W. Lee, et al., 2014). Nevertheless, many real-life decisions are made in the absence of external, value-based outcomes, where agents need to form beliefs based on other sources of information. Partially addressing this issue, studies on perceptual decision making have explored how people make choices based only on sensory evidence, in the absence of outcomes (Hanks & Summerfield, 2017; Heekeren, Marrett, & Ungerleider, 2008). These studies focus on attention processes and perceptual uncertainty, as sensory inputs are characterized by ambiguity or noise, and have highlighted the roles played by hippocampus and mPFC in assessing sensory predictability and the subsequent choice confidence (Bang & Fleming, 2018; Harrison, et al., 2006; Rahnev, Nee, Riddle, Larson, & D’Esposito, 2016; Strange, et al., 2005). Instead, here we aimed at investigating the neural dynamics responsible for the update of action-outcome contingencies in the presence and absence of outcomes. Both our two multi-option tasks, with and without immediate value-based feedback, were characterized by simple sets of rules, categorical and discrete evidence (i.e. the three feedback values or the three bead colors), and easy-to-compute distributions of probabilities, reducing stimulus uncertainty as well as second-order uncertainty (Bang & Fleming, 2018; Fleming & Daw, 2017).

DCM analysis indicates that the directional influence among brain regions encoding decision confidence changed across environments. In the presence of explicit outcomes, mPFC and amygdala drove other regions during confidence encoding; but in the absence of these outcomes, the hippocampus showed directional influence towards mPFC and amygdala. Confidence estimations in the bead task (no outcome) relied on each colored bead in a sequence as discrete evidence, where a sequence of beads of the same color signaled that the environment was likely going through a stable phase and therefore decision confidence could increase during this phase. Therefore, we speculate that the hippocampus was engaged first in order to monitor the predictability of visual stimuli and signal the reduction in entropy of the task environment (Harrison, et al., 2006; Rigoli, et al., 2019; Strange, et al., 2005), as belief-confirming evidence was accumulating. Subsequently, the mPFC and amygdala received this information from the hippocampus (Gluth, Sommer, Rieskamp, & Buchel, 2015) and were respectively engaged to increase the estimated confidence (Bang & Fleming, 2018; Matsumoto & Tanaka, 2004; Yoshida & Ishii, 2006) and to assign a value to each selected action (Bechara, et al., 1999; Dolan, 2007; Jung, et al., 2018). Differently, in the card task where stochastic numeric outcomes were available, evidence accumulation was based on expected values. Thus, we speculate that the mPFC (De Martino, et al., 2013; Koechlin & Hyafil, 2007) and amygdala (Bechara, et al., 1999; Dolan, 2007; Jung, et al., 2018) became the driving force in calculating these value-based signals and assigning them to the available choices. This information was then passed to the hippocampus, to monitor the stability or entropy of the environment.

In contrast, we found that the AIC drove the uncertainty-building network and exerted its directed influence over dACC and dlPFC, regardless of the presence of immediate value-based outcomes. The AIC-dACC-dlPFC circuit has been consistently implicated in various cognitive processes (e.g. attention, response selection, cognitive control), leading to the hypothesis that they are part of a ubiquitous salience network (Seeley, et al., 2007; Uddin, 2015). This network also monitors threats to homeostasis (Gu & FitzGerald, 2014), as in the case of information conflicting with established beliefs, as for instance in reward (Behrens, et al., 2007; Gu, Wang, et al., 2015; Rushworth & Behrens, 2008; Seymour, et al., 2004) or state (Glascher, et al., 2010; S. W. Lee, et al., 2014) prediction errors, which are essential to determine confidence reduction/uncertainty building across our tasks. In this sense, our finding of the AIC driving the uncertainty network regardless of outcome availability is consistent with its role in salience processing and belief updating (Gu & FitzGerald, 2014; Uddin, 2015).

It is important to highlight a few limitations associated with the interpretation of the behavior recorded in the two tasks. Despite the described core of similarities, the two tasks differ in some key aspects: first, the chronicle of the latest 5 outcomes is only externalized in the bead task; second, the bead task is observation-based, whereas the card task is action-based; finally, the card task presents an increased exploration cost in comparison with the bead task. The high level of accuracy associated with the Bayesian learner model provides indications about the impact these limitations have on the interpretation of our findings. In both tasks the update of choice-confidence and uncertainty are based on the observation of the latest available data, therefore reducing the impact of the chronicle externalization. Furthermore, this feature also implies both tasks are treated as if they were observation based. In the card task the observations are limited to the choice selections performed in the previous trials, but the information provided either confirms or conflicts with existing belief by the same quantity in both tasks, due to the organization of evidence into three distinct categories. Finally, despite the higher cost of exploration in the card task, in comparison with the bead task, participants were found to explore more in the card task, as highlighted in the lower estimated σ values and associated faster pace of update. All considered, we suggest that these limitations call for further investigations, but they do not have a major impact on the interpretation of the key findings described in this study.

Taken together, our findings confirm the importance of multi-level investigations across the Marrian tri-level of analysis and shed a new light on pervasive computational and neural mechanisms underlying belief formation and update. These results represent also an important step to inform future investigations into the breakdown of belief update processes, such as those observed in addiction (Gowin, Mackey, & Paulus, 2013; Ognibene, Fiore, & Gu, 2019; Verdejo-Garcia, Chong, Stout, Yucel, & London, 2018), mood disorders (Bishop & Gagne, 2018; Huys, Daw, & Dayan, 2015), as well as across several psychiatric disorders (Hoven, et al., 2019).

## Acknowledgments

We thank Prof. Karl Friston for his comments and kind suggestions in shaping this manuscript. We thank Jennifer Jung for her help in setting up the code for both tasks (Psychtoolbox library in Matlab), Ann-Cathrin V. Guertler, Chandana C. Tatineni, and Ju-Chi Yu, for their help in collecting the data. VGF is funded by the Mental Illness Research, Education, and Clinical Center (MIRECC VISN 2) at the James J. Peter Veterans Affairs Medical Center, Bronx, NY. XG is supported by National Institute of Drug Abuse (R01DA043695, R21DA049243) https://www.drugabuse.gov/, and the National Institute of Mental Health (R21MH120789) https://www.nimh.nih.gov/. The funders had no role in study design, data collection and analysis, decision to publish, or preparation of the manuscript. The authors have declared that no competing interests exist.

